# Model-Enabled Knowledge Transfer across cell lines, culture scales and conditions

**DOI:** 10.64898/2025.11.30.691385

**Authors:** Luxi Yu, Antonio del Rio Chanona, Cleo Kontoravdi

**Affiliations:** Department of Chemical Engineering, South Kensington campus, Imperial College London, London SW7 2AZ, United Kingdom

**Keywords:** Bioprocess Modelling, State Estimation, Ensemble Kalman Filter, Knowledge Transfer

## Abstract

Mechanistic models are central to quantitative understanding and optimisation of Chinese hamster ovary (CHO) cell culture processes, but their utility is often restricted by parameter sets calibrated for specific cell lines, scales, or operating conditions. In this study, we present the application of the Ensemble Kalman Filter (EnKF) to bioprocessing, introducing an ensemble-based framework for dual state and parameter estimation that enables mechanistic model adaptation across distinct systems. The EnKF recursively assimilates process measurements to update uncertain kinetic parameters and predict system states, allowing a model developed for one system to be transferred to a new one without reparametrisation and using only a single experimental dataset. The evolving parameter ensembles provide a time-resolved sensitivity analysis that identifies which parameters have dominant influence under new process conditions and when their effects become significant. The framework was evaluated across six CHO cell experimental datasets differing in scale, cell line, temperature, and feeding strategy, demonstrating accurate reconstruction of system dynamics and progressive improvement in long-term predictions as new data became available. By maintaining full mechanistic transparency while flexibly adapting to new data, the EnKF offers a practical route for knowledge transfer across systems, strengthening the role of mechanistic modelling in data-informed bioprocess understanding and control.

## 1 Introduction

Effective knowledge transfer across bioprocessing systems is essential for reducing process development timelines, adapting platforms to new cell lines and products, facilitating scale-up and optimising production [1, 2]. However, inherent biological variability and system-specific differences in metabolism, growth dynamics, and, consequently, nutrient uptake rates often limit the transferability of process insights across systems [3]. In the absence of a structured methodology for adopting prior knowledge, process development continues to rely on extensive experimentation, leading to increased cost and longer timelines [4].

Mathematical models offer a systematic framework for understanding and predicting bioprocess behaviour through quantitative description of system dynamics [5]. Mechanistic models, in particular, are widely employed due to their ability to explain biological and physicochemical behaviours based on first principles [6, 7]. While these models can provide valuable insight into cell behaviour, their predictive performance is often constrained by system-specific parameterisation [8]. When applied to different bioreactor scales, cell lines, or operating regimes, mechanistic models typically require extensive reparametrisation to maintain accuracy [9].

To overcome these challenges, recent efforts have focused on data-driven and hybrid modelling approaches designed to improve model transferability. Stand-alone data-driven methods, such as transfer learning using Artificial Neural Networks (ANNs) or multi-fidelity Gaussian process models, are flexible and often perform well in data-scarce regimes [10–13]. However, they typically lack mechanistic interpretability. Hybrid models improve generalizability while partially retaining interpretability by combining mechanistic structure with data-driven components, as demonstrated in several recent studies [8, 14–18]. Yet in many cases, the mechanistic backbone is reduced to mass balances, with core biological behaviours such as metabolic fluxes and growth kinetics captured by black-box surrogates [14, 19], reducing the model’s capacity to provide mechanistic insight. In other hybrid implementations, system identity is encoded via one-hot vectors [20] or embedding layers [21], allowing the model to distinguish between systems based on historical data. However, they do not directly identify the underlying physiological traits responsible for these differences. Moreover, although hybrid models are less data-intensive than fully data-driven approaches, they still require multiple datasets to train their data-driven components. This presents a challenge in early-stage process development, where reliable predictive models are most needed but data is scarce [22]. In addition, the absence of explicit uncertainty handling increases the risk of overfitting, particularly when datasets are limited or noisy [23].

A fundamental limitation shared across mechanistic, data-driven, and hybrid models is their reliance on fixed parameter values across process duration. Once parametrised, these models operate deterministically and are typically applied offline, limiting their ability to adapt to deviations in system behaviour caused by parameter uncertainty, measurement noise, or biological variability. This motivates the need for dynamic modelling frameworks that incorporate real-time measurements to update both states and parameters while retaining mechanistic structure. In this context, state estimation algorithms offer a practical alternative.

Various state estimation techniques have been applied to bioprocess applications, each with distinct advantages and limitations. The Extended Kalman Filter (EKF) remains one of the most widely used methods due to its easy implementation and computational efficiency [24–30]. EKF linearises the process model and is generally suitable for mildly nonlinear systems [31]. However, its accuracy deteriorates when applied to strongly nonlinear and discontinuous processes, which are common in mammalian cell cultures [32]. The Unscented Kalman Filter (UKF) addresses nonlinearities more effectively than the EKF by propagating sigma points through the nonlinear model, with successful applications reported in the bioprocessing literature [33–40]. While more robust than the EKF, the performance of UKF is sensitive to the choice of hyperparameters, often requiring careful tuning to ensure stability and accuracy. Particle Filter (PF) has been applied to many complex bioprocess systems that exhibit highly nonlinear dynamics and non-Gaussian uncertainty [41–45], at the expense of higher computational demand [46]. Apart from recursive methods, Moving Horizon Estimation (MHE) formulates the state estimation problem as a constrained optimisation over a finite time horizon [47–49], explicitly accounts for model constraints and past measurements [50–52]. However, the need to solve an optimisation problem introduces substantial computational overhead, which can limit its use for real-time applications.

The Ensemble Kalman Filter (EnKF) offers a promising solution to many of the challenges associated with state estimation techniques of bioprocessing systems. Originally developed for large-scale systems in geosciences and reservoir modelling [53], EnKF combines the computational efficiency of Kalman-based methods with the flexibility of ensemble sampling to handle nonlinearity and uncertainty. Unlike the EKF and UKF, it requires no Jacobian computation or sigma point selection, avoids the high computational burden of PF, and the need to solve an optimisation problem in MHE. By propagating an ensemble of model realisations, the EnKF naturally captures the evolution of uncertainty in both states and parameters, without requiring the explicit computation or storage of process covariance matrices [54]. This ensemble-based formulation offers a favourable balance among estimation accuracy, ease of implementation, and computational efficiency.

We have previously applied the EnKF for soft sensing of unmeasured intracellular states in CHO cell cultures [55]. In this study, we extend its application beyond state estimation to the recursive updating of uncertain kinetic parameters throughout the duration of the culture. By analysing the temporal evolution of parameter ensembles, we could gain insights into dynamic changes in cellular behaviour that would otherwise be masked by static calibration. As the parameter trajectories are continuously refined based on incoming measurements, this approach preserves the mechanistic structure while adapting flexibly to new conditions without the need for reparametrisation. This capability supports knowledge transfer by allowing a model developed for one specific system to be applied to different cell lines, scales, or operating conditions using only a single experimental dataset. Moreover, the evolving parameter trajectories enable a form of dynamic sensitivity analysis, where the propagation of ensemble uncertainty highlights which parameters are most influential under the new system conditions.

## 2 Materials and Methods

### 2.1 Mechanistic Model and Experimental Data

#### 2.1.1 Base Mechanistic Model and Calibration Dataset

The mechanistic model used for EnKF implementation was adapted from the cell culture framework described by Kotidis *et al*. (2019) [56], with model equations provided in Supplementary Information A. This reduced model captures key extracellular dynamics, including cell growth and death, metabolism, and antibody synthesis, while excluding intracellular modules related to nucleotide sugar metabolism and glycosylation. The experimental data for this mechanistic model calibration were obtained from fed-batch cultures of an IgG-producing CHO-T127 cell line. The culture were maintained in CD CHO medium (Life Technologies, Paisley, UK) at 36.5°C with 5% CO_2_ and agitated at 150 rpm. Cells were passaged every 3-4 days, with 50 *µ*M of methionine sulfoximine supplemented during the initial two passages after revival. Cultures were grown in 500 mL vented Erlenmeyer flasks (Corning, Amsterdam, The Netherlands) with a 100 mL working volume and an initial seeding density of 2 × 10^5^ cells mL^-1^. 10% (v/v) CD EfficientFeed C™ C AGT™ (Life Technologies, Paisley, UK) was added every alternate day starting from day 2 [56].

To investigate whether the EnKF can adapt a fixed mechanistic model beyond its original parametrisation, we applied the method to datasets from diverse bioprocess conditions. These included changes in bioreactor scale, temperature, host cell line, and feeding strategy, as summarised in Table 1. The structure of the model remained unchanged across all cases, with only model parameter updated recursively during the culture.

**Table 1:** Summary of experimental datasets. Only the feed concentration of nutrients considered in the model is reported.

| # | Cell Line | Bioreactor |  |  | Feed Concentration (mM) |  |  | Ref |
| --- | --- | --- | --- | --- | --- | --- | --- | --- |
|  |  | Type | Volume (mL) | Temp (°C) | Glc | Glu | Asn |  |
| 1 | A | Shake flask | 100 | 36.5 | 144.4 | 12.2 | 27.0 | [56] |
| 2 | A | Bioreactor | 900 | 36.5 | 144.4 | 12.2 | 27.0 | [57] |
| 3* | A | Bioreactor | 900 | 32 | 144.4 | 12.2 | 27.0 | [57] |
| 4 | B | Shake flask | 50 | 36.5 | 144.4 | 12.2 | 27.0 | [58] |
| 5 | B | Shake flask | 50 | 36.5 | 144.4 | 21.4 | 73.6 | [58] |
| 6 | B | Shake flask | 50 | 36.5 | 202.2 | 29.9 | 103.1 | [58] |

#### 2.1.2 Same Cell Line: Scale and Temperature Variation

The first three datasets correspond to the same CHO-T127 cell line (Cell Line A). Dataset 1 represents the original shake-flask culture used for model calibration, whereas Datasets 2 and 3 were conducted in 1.5 L stirred-tank bioreactors (DASGIP Technology, Jülich, Germany) to evaluate model transferability across scales and temperature conditions. Both bioreactor datasets were seeded at 8 × 10^5^ cells mL^-1^ with a working volume of 0.9 L and maintained at pH 6.9 *±* 0.1, 5 % CO_2_, 150 rpm agitation, and active antifoam control. In Dataset 2, the culture temperature was held constant at 36.5 °C, while in Dataset 3, a temperature downshift from 36.5 °C to 32 °C was introduced on day 6. The bioreactor cultures followed the same feeding schedule as the shake-flask experiment, with 10 % (v/v) of Feed C added on even-numbered days starting on day 2, and were further supplemented with 10 % (v/v) of a proprietary nutrient Feed X (MedImmune, Cambridge, UK) [57].

#### 2.1.3 Different Cell Line: Feeding Strategy Variation

To evaluate the capability of EnKF to support cross-cell line knowledge transfer, three additional datasets based on CHO-GS46 (Cell Line B) were tested, each employing a distinct feeding strategy. Cultures were incubated in 250 mL Erlenmeyer flasks (Corning, Amsterdam, The Netherlands) with a 50 mL working volume, incubated at 36.5°C, 140 rpm, and 8% CO_2_. Feeding was performed every other day using a tenfold-concentrated nutrient supplement at 10% (v/v). Dataset 4 employed the same commercial feed used in Cell Line A shake flask culture. Datasets 5 and 6 used in-house formulations Feed U and Feed U plus 40%, developed from the batch culture nutrient consumption profiles of Cell Line B. Amino acid and glucose concentrations were scaled to sustain a threefold increase in integrated viable cell concentration, with the latter formulation containing 40 % higher nutrient levels except for tyrosine, which was limited by solubility. While the feeds contained a broader range of components, only the nutrients represented in the model are listed in Table 1 [58].

#### 2.1.4 Analytical Methods

Cell concentration and viability were measured either manually using a Neubauer haemocytometer with trypan blue exclusion for shake flask samples or with the automated ViCell® system (Beckman Coulter, CA) for bioreactor cultures. Antibody titre in shake flask samples was quantified using the BLItz® system with Dip and Read™ Protein A biosensors (Pall ForteBio, Portsmouth, UK), whereas in the bioreactor studies, secreted mAb concentration was determined by Protein A affinity chromatography. Extracellular glucose, glutamine, glutamate, lactate, and ammonia concentrations were measured using the BioProfile 400 analyser (NOVA Biomedical, MA) for shake flask experiments, while bioreactor samples were analysed using the YSI Bioprofiler 800 (NOVA Biomedical, MA). Extracellular amino acid concentrations, including asparagine, were quantified by high-performance or ultra-performance liquid chromatography (HPLC/UPLC, Waters, Hertfordshire, UK) using the AccQ-Tag™ kit according to the manufacturer’s protocol.

### 2.2 EnKF Dual State and Parameter Estimation

Figure 1 illustrates the structure and information flow of the dual EnKF implementation. At each update step, ensembles of parameters and states, initially sampled from prior distributions, are propagated through the mechanistic model. As new measurements become available, the parameter and state ensembles are sequentially updated, reducing their spread and thereby the associated uncertainty. The narrowing of these distributions reflects convergence towards the true system dynamics, enabling continuous refinement of both parameters and states throughout the culture.

**Figure 1.**
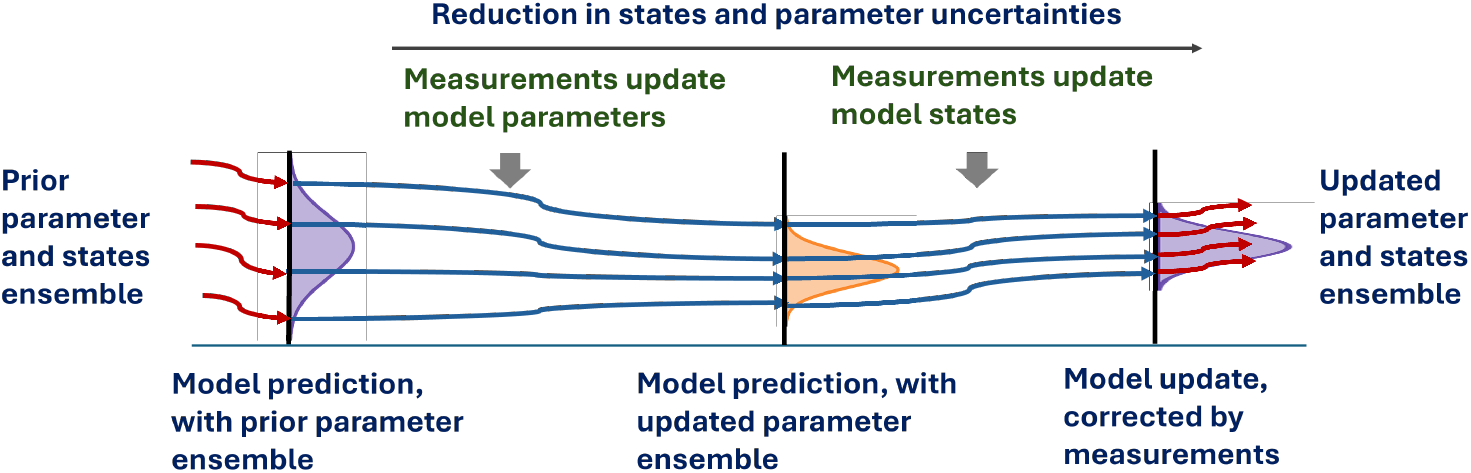
Schematic of EnKF dual state and parameter estimation.

**Figure 2.**
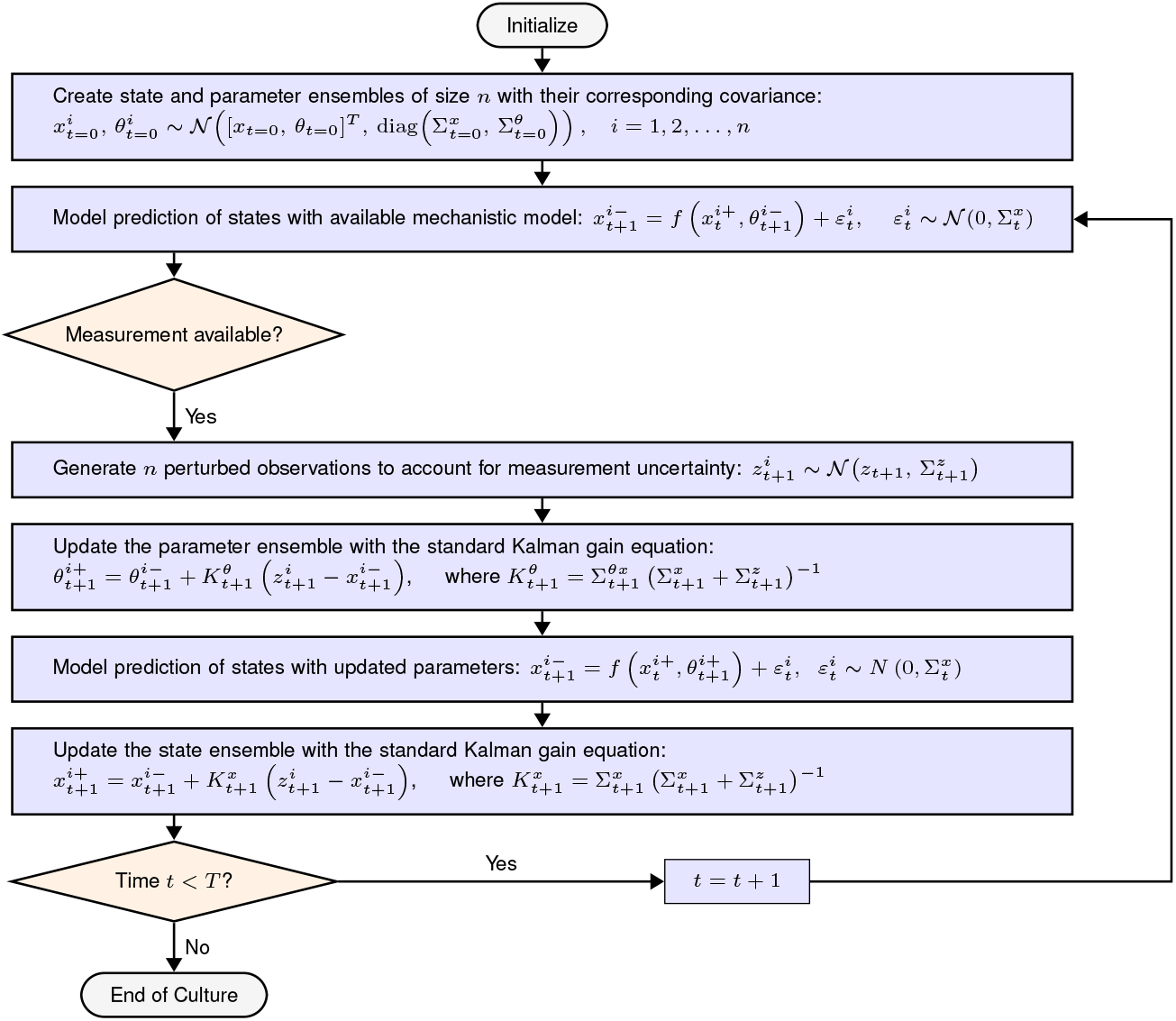
EnKF dual state and parameter estimation workflow.

The ensemble is initialised at *t* = 0 with *n* members, each obtained by independently sampling the state and parameter vectors from Gaussian distributions centred at their prior nominal values, thereby generating *n* distinct realisations that collectively represent the prior uncertainty:

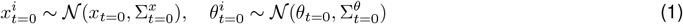

where 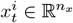 is the state vector and 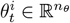 is the parameter vector for the *i*th ensemble member, and *i* 1,2, …, *n*. Superscripts − and + denote variables before and after the measurement update, respectively. The covariance matrices 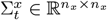 and 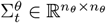 represent the process and parameter uncertainty at time *t*.

The mechanistic model is based on Kotidis *et al*. (2019), with published parameter values used as the prior mean. For each dataset, the initial ensemble covariance reflects dataset-specific uncertainty in parameters and states. The ensemble of prior states evolves through the model using the prior parameter ensemble, with Gaussian-distributed process noise added to account for model uncertainty:

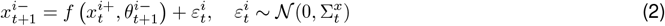

Due to PAT constraints, on-line measurements are sparse. The mechanistic model is numerically integrated with a fine internal time step to capture system dynamics continuously, while measurement updates occur only when experimental data become available, typically once per day in fed-batch cultures. When off-line measurements are obtained, the reported experimental mean is used as the central value, and an observation ensemble is generated to reflect the associated measurement uncertainty [53]:

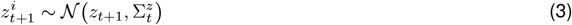

where 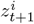 denotes the *i*th perturbed measurement and 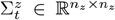 is the measurement covariance matrix, representing the uncertainty and potential correlations among measured process variables. The experimental error bar can be used as a practical reference for measurement covariance matrix, accounting sampling and instrument uncertainty. A lower bound is enforced to prevent sampling negative concentrations. The measurements are then used to correct the model parameter ensemble. The parameter update follows the standard Kalman correction:

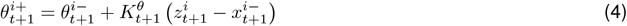

The update is scaled by the parameter Kalman gain 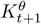, which determines how strongly each parameter is adjusted based on the current mismatch between model prediction and measurement.

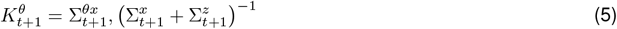

where 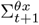 is the cross-covariance between the parameter and predicted state ensembles. The updated parameter ensemble is then used to re-predict the states:

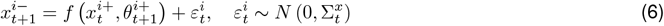

The predicted states are subsequently corrected using the states Kalman gain 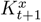, which balances the confidence in the model prediction against the reliability of the experimental data.

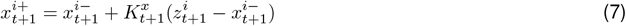

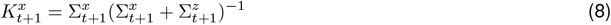

In this work, a dual estimation structure was adopted instead of a fully joint formulation in which states and parameters are concatenated into a single augmented vector. In the state update, stochastic process noise is explicitly introduced at each propagation step to capture model uncertainty, effectively inflating the covariance matrix dynamically. For parameters, no artificial noise is injected, ensuring that changes in ensemble spread reflect genuine information gain from experimental data rather than covariance inflation. By maintaining separate ensembles, state propagation remains numerically stable while parameter evolution can retain a relatively large initial covariance to permit adaptive learning. If a large initial covariance were assigned to the augmented state-parameter vector, the resulting integration would become computationally unstable and inefficient due to the need for repeated numerical integration with overly dispersed trajectories. This separation therefore facilitates both stable computation and biologically interpretable parameter trajectories.

### 2.3 EnKF Uncertainty Quantification

The mechanistic model consists of eight state variables and twenty-four kinetic parameters. To reflect the scale and complexity of the system, an ensemble size of 50 was used for the three Cell Line A datasets. For the Cell Line B datasets, a larger ensemble of 75 was selected to account for their greater deviation from the base model and the increased process uncertainty. The accuracy of state and parameter estimation further depends on how uncertainty is represented and propagated through the model. Three sources of uncertainty were specified, namely measurement noise, process noise, and parameter covariance.

#### 2.3.1 Measurement Noise and Process Noise Tuning

Measurement noise 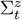 represents observation uncertainty arising from instrumentation, sampling, and handling variability [54, 59]. Initial estimates were based on the standard deviations inferred from experimental error bars. Biological replicates were treated with caution to avoid overestimating measurement noise, which may also reflect process variability. Technical replicates are preferred where available to isolate instrument-specific error.

Process noise 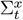 accounts for model-structure limitations, unmodelled dynamics, and residual process disturbances. It was initially estimated from model-measurement discrepancies and tuned in conjunction with 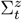 to regulate the Kalman gain. The ratio between process and measurement noise influences how strongly the filter relies on model predictions versus observed data. This ratio was iteratively adjusted for each dataset to reflect data quality, confidence in model structure, and expected deviations from calibration conditions.

In this work, the model is applied to conditions beyond its original calibration, including different bioreactor scales, cell lines, and feeding strategies. To reflect reduced model confidence under these conditions, the process noise 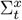 is set relatively high compared to 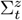, ensuring the filter places more weight on measurement data during adaptation. Final tuned values for 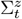 and 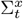 are provided in the Supplementary Information B.

#### 2.3.2 Parameter Spread and Dynamic Sensitivity Analysis

A common initial parameter covariance matrix 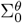 was applied across all datasets, yielding identical initial spreads for each parameter and allowing consistent interpretation of parameter sensitivity under different process conditions. The initial ensemble standard deviations were selected to be high but physiologically plausible, preventing early ensemble collapse and enabling flexibility for parameter adaptation. Unlike traditional model fitting, which yields static confidence intervals, the EnKF framework updates the ensemble spread dynamically as experimental data are assimilated. As the culture progresses and more measurements are incorporated, parameter uncertainty typically decreases. To quantify this reduction, the relative decrease in ensemble standard deviation for each parameter over the course of the culture was defined as:

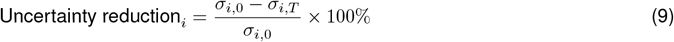

where *σ*_*i*,*t*_ denotes the ensemble standard deviation of the *i*th parameter at time *t*. This metric serves as a proxy for dynamic sensitivity analysis, indicating which parameters are most informed by the available data under new process conditions. Parameters showing substantial uncertainty reduction are strongly constrained by the data, while those with little or no reduction may be weakly identifiable or insensitive under the given operating regime. Full numerical values for the initial ensemble spread and percentage reduction across parameters are reported in the Supplementary Information C and D.

## 3 Results and Discussion

### 3.1 Model Transfer from Shake Flask to Bioreactor and Temperature Downshift

#### 3.1.1 State Estimation Across Scale and Temperature Shift

The shake flask culture (Dataset 1, grey) provided the base mechanistic model, which was then adapted to the bioreactor operated at 36.5 °C (Dataset 2, purple) and 32 °C (Dataset 3, teal). The resulting EnKF predictions for all three conditions are shown in Figure 3. The filter facilitates recursive parameter and state updates throughout the culture without model reparametrisation.

**Figure 3.**
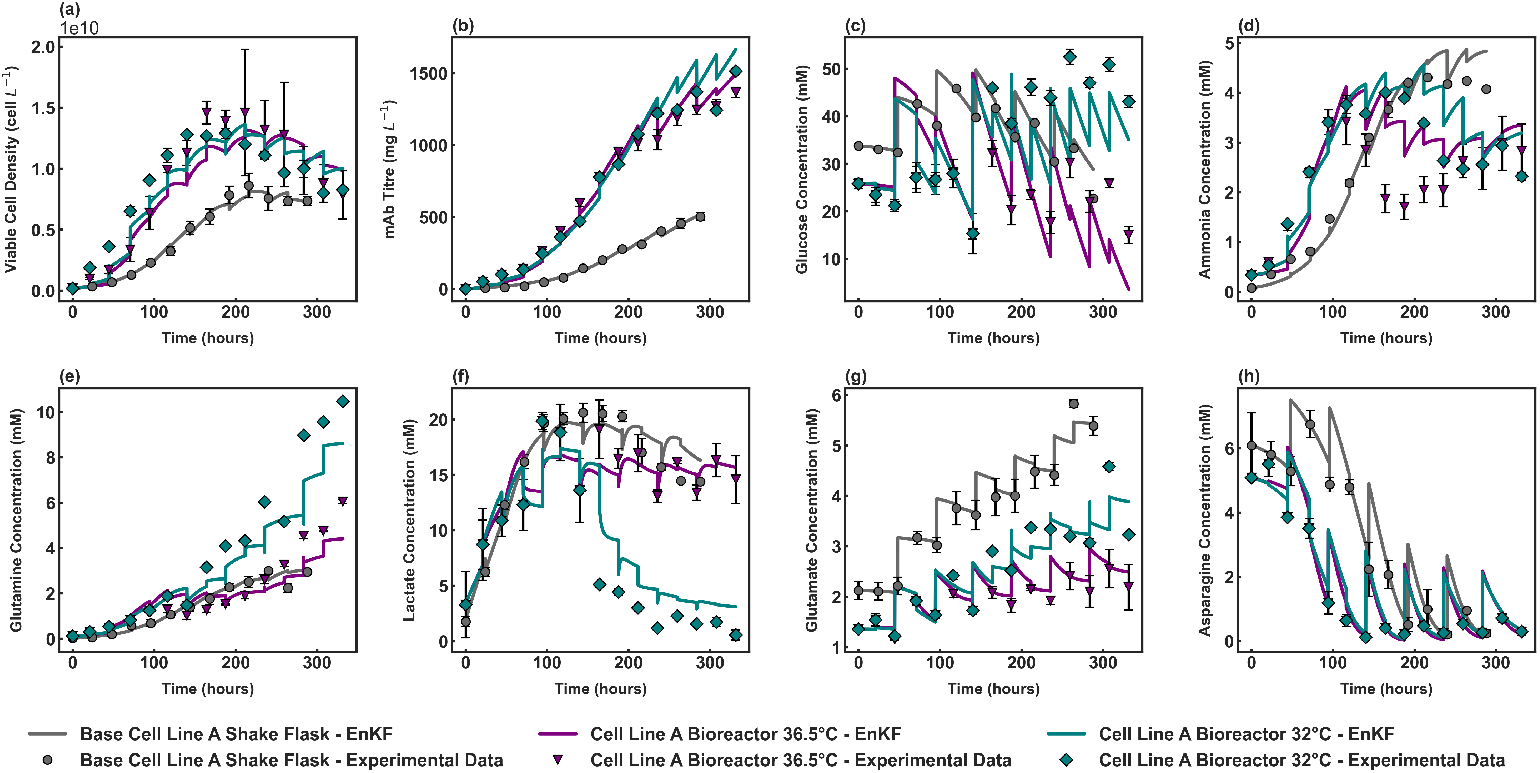
EnKF-based state estimation model transfer across scale and temperature downshift for Cell Line A cultures.

**Figure 4.**
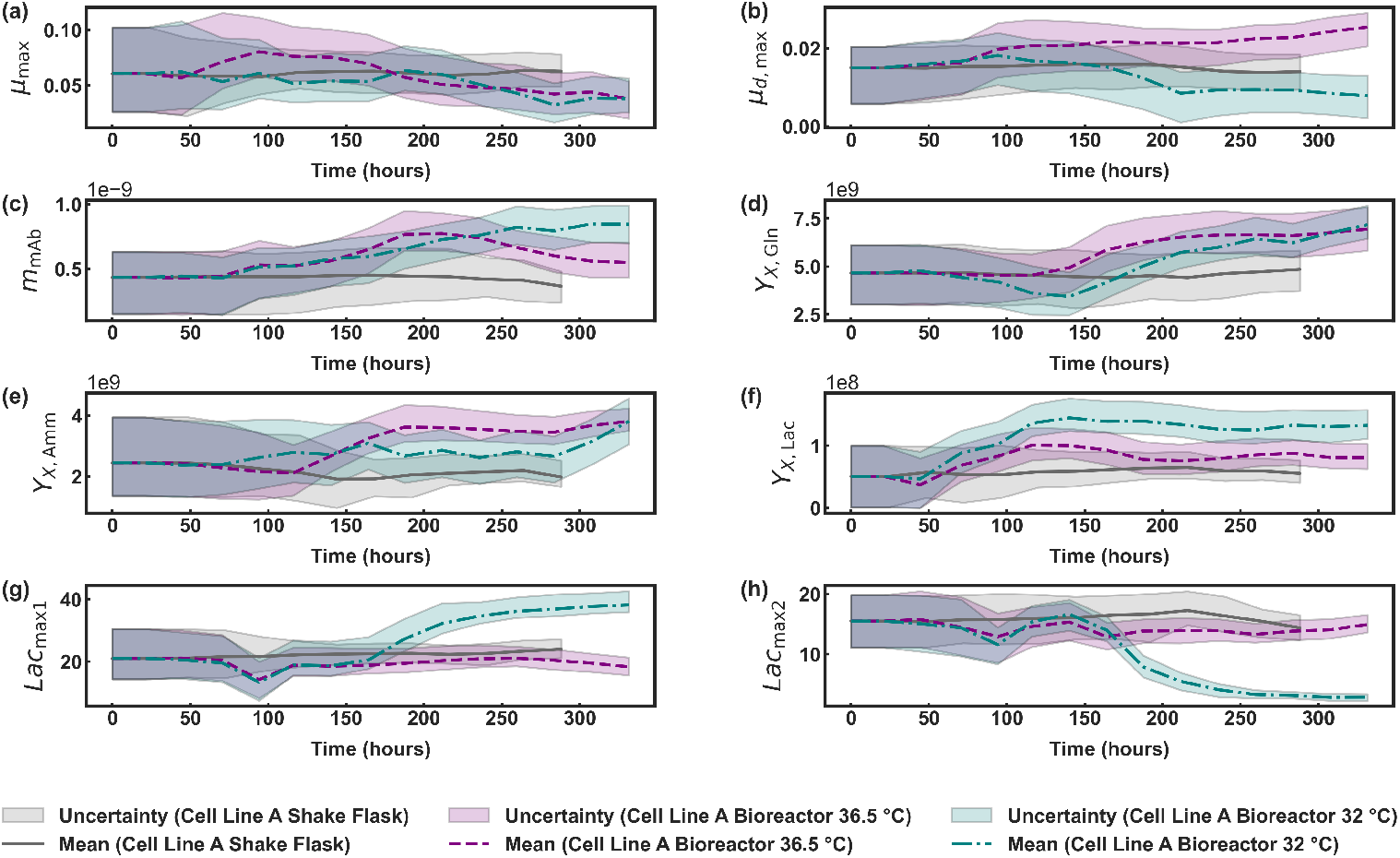
Time evolution of estimated model parameters across scale and temperature downshift for Cell Line A cultures.

The higher cell growth and mAb productivity observed at bioreactor scale were both well captured. The EnKF accurately reproduced the overall glucose consumption trends across all three conditions. A slight misalignment was observed under the 32°C condition, which may be attributed to uncertainty in the composition of the proprietary Feed X added in addition to the commercial Feed C. Although Feed X was modelled as part of the nutrient input, its compositional uncertainty was not explicitly represented in the EnKF, meaning its variability was not propagated through the ensemble. Ammonia accumulation plateaued earlier at bioreactor scale compared to shake flask, with the EnKF recovering the general trajectory. Importantly, the filter did not track the ammonia dip between days 6 and 10 for Bioreactor 36.5°C dataset, indicating that it did not overfit to what may represent measurement noise rather than a true metabolic shift.

Following the temperature downshift, glutamine concentrations in the Bioreactor 32°C dataset were substantially elevated, and this was reliably captures by the filter. A distinct shift in lactate metabolism was also observed in response to the temperature downshift, with net lactate consumption occurring after day 6, in contrast to the accumulation observed in shake flask and 36.5°C bioreactor conditions. The lactate consumption trend is successfully captured but the end concentration is slightly lower than that described by the EnKF.Glutamate concentrations varied across datasets, yet the predicted trajectories followed the observed profiles across all three conditions. Asparagine dynamics exhibited clear scale dependence, with the EnKF accurately describing the rate and extent of depletion.

#### 3.1.2 Parameter Evolution for Knowledge Transfer Across Scale and Temperature Shift

##### Cell growth and productivity

Parameter values remain relatively constant across culture duration in the shake flask experiments. This is expected given that this dataset was used for model parametrisation and the initial ensemble adequately captures baseline behaviour. The trajectory of the maximum specific growth rate *µ*_*max*_ and death rate *µ*_*d*,*max*_ aligns well with the observed viable cell density profile shown in Figure 3 (a). In the bioreactor experiments conducted at 36.5°C, *µ*_*max*_ is increased during early exponential growth, reflecting the higher viable cell densities achieved in this controlled environment with stable pH and dissolved oxygen tension. Interestingly, the trajectory of *µ*_*d*,*max*_ shows higher rate of cell death in bioreactors compared to shake flask cultures, potentially because the higher viable cell density achieved. In the 32°C bioreactor, however, a clear reduction in *µ*_*d*,*max*_ is observed following the temperature downshift, indicating delayed and slower cell death. This behaviour is consistent with the expected benefits of mild hypothermia in CHO cultures, which has been shown to prolong viability and extend the productive phase [57, 60]. These advantages are also evident in the trajectory of the titre production parameter *m*_*mAb*_, which reaches its highest values under 32°C, reflecting enhanced specific productivity with temperature downshift.

##### Scale-dependent glutamine utilization

The yield of biomass from glutamine *Y*_*x*,*Gln*_ remains relatively constant with time in the shake flasks but increases in both bioreactor cultures from day 6 onwards, reflecting enhanced glutamine utilisation for cell growth at larger scale and in a controlled environment. In GS-CHO cell lines, glutamine is synthesised from glutamate and ammonia via an ATP-dependent glutamine synthetase reaction [61]. As ATP is primarily generated through oxidative phosphorylation, the rate of this conversion is sensitive to oxygen availability [62]. Improved oxygen transfer in bioreactors supports sustained ATP production, thereby enabling more efficient glutamine synthesis from ammonia. This mechanistic interpretation is further supported by a parallel increase in the yield of biomass from ammonia *Y*_*x*,*Amm*_. Although glutamine concentration profiles appear similar in the shake flask and 36.5°C bioreactor until day 12 as seen in Figure 3 (e), the dynamic trajectory of *Y*_*x*,*Gln*_ inferred by the EnKF reveals scale-dependent effects as early as day 6.

##### Lactate consumption induced by temperature shifts

The yield of biomass on lactate *Y*_*X*,*Lac*_ reaches its highest values in the 32°C bioreactor, suggesting increased utilisation of lactate as an energy source following the temperature downshift. Scaling up from shake flask to bioreactor at 36.5°C does not substantially alter lactate accumulation, and this is reflected in the relatively stable trajectories of *Lac*_*max*1_ and *Lac*_*max*2_, which remain close to the base model.

These parameters represent kinetic constants that regulate the activation of lactate consumption pathways. In the 32°C cultures, *Lac*_*max*1_ increases during the later stages indicating lactate accumulation, while *Lac*_*max*2_ decreases sharply after the downshift due to net lactate consumption. The parameter trajectories provide evidence for the stronger influence of *Lac*_*max*2_ on net lactate consumption, reflecting the bidirectional nature of lactate metabolism.

### 3.2 Model Transfer from Cell-line A to B under Different Feeds

#### 3.2.1 State Estimation Across Cell-lines and Feeding Conditions

The base model, originally parametrised using Cell Line A shake flask data supplemented with Feed C, was transferred to the Cell Line B. Three feeding strategies were evaluated, Feed C (Dataset 4, orange), Feed U (Dataset 5, green), and Feed U plus 40% (Dataset 6, blue), with all studies conducted in shake flasks. Despite differences in cell line and feed compositions, the EnKF successfully adapted the base model to capture cell growth, productivity, and metabolite dynamics for all three cultures.

Distinct growth and death profiles were observed across the feeding strategies. Compared to Feed C, Feed U and Feed U plus 40% both resulted in a steeper decline in viable cell density following peak growth. These differences were accurately captured by the EnKF, even though all predictions were based on the same underlying model structure. The mAb titre profiles were also well reproduced, with Feed C yielding the highest final titre of approximately 2.5 g L^-1^. This represents a nearly four-fold increase compared to the baseline model calibrated on Cell Line A, yet the EnKF successfully captured this shift, demonstrating transferability across systems with markedly different productivity levels.

The higher concentrations of glucose and asparagine in Feed U and Feed U plus 40% resulted in different metabolite profiles compared to Feed C. In particular, excess asparagine feeding led to substantial ammonia accumulation, with concentrations exceeding 20mM by the end of the culture under Feed U plus 40%. While the EnKF described the general trend well, some underprediction was observed during the final stage in this condition. During filter tuning, it was found that a higher ammonia measurement noise was required to stabilise the EnKF in the Feed U plus 40% case. Reducing the measurement noise resulted in a closer fit to the ammonia data but introduced instability at later time points, as the filter was forced too closely towards the measurements and propagated aggressive corrections into correlated states, particularly glutamine, glutamate and asparagine. The slight mismatch observed is therefore expected, given the greater uncertainty in ammonia measurements at high concentrations. Nonetheless, the EnKF well represented the broader dynamics of nitrogen metabolism across conditions, including elevated glutamine and glutamate concentrations in both Feed U and Feed U plus 40%.

Lactate profiles further highlight the flexibility of the EnKF framework. The base model parametrised on the Cell Line A shake flask dataset exhibited net lactate accumulation under Feed C, as shown in Figure 3 (f). In contrast, Cell Line B culture under the same scale and feeding strategy showed net lactate consumption, indicating this behaviour is cell line-specific. Feed U and Feed U plus 40% resulted in even stronger lactate consumption, with near-complete depletion by the end of the culture in the latter case. These distinct profiles were accurately described using the same underlying mechanistic model, made possible by the dynamic evolution of the parameter ensemble. Such flexibility would not be attainable under a fixed-parameter model calibration.

#### 3.2.2 Parameter Evolution for Knowledge Transfer Across Cell-lines and Feeding Conditions

##### Cell growth and productivity

The trajectory of the maximum specific death rate *µ*_*d*,*max*_ provides a clear distinction between the commercial Feed C and the in-house formulations, Feed U and Feed U plus 40%. Ensemble-based uncertainty narrows after the exponential growth phase, reflecting the higher sensitivity of *µ*_*d*,*max*_ during later stages of cell culture. In Cell Line B supplemented with Feed C, *µ*_*d*,max_ remains relatively constant throughout the culture, within the uncertainty of the parameter estimate. In contrast, Feed U shows an increase in the mean death rate during the stationary phase, which is expected given its limited nutrient composition compared to the commercial feed and the associated rise in ammonia levels. Feed U plus 40 % exhibits an even higher *µ*_*d*,max_, consistent with the pronounced ammonia accumulation observed in this condition and its strong correlation with increased cell death, as illustrated in Figure 5 (a). Among the three feeding strategies, Feed C supports the highest titre yields, while the two in-house formulations result in comparable but lower final titres, as shown in Figure 5 (b). This suggests that improved cell viability under Feed C is accompanied by higher non-growth-associated productivity *m*_*mAb*_, contributing to enhanced overall production.

**Figure 5.**
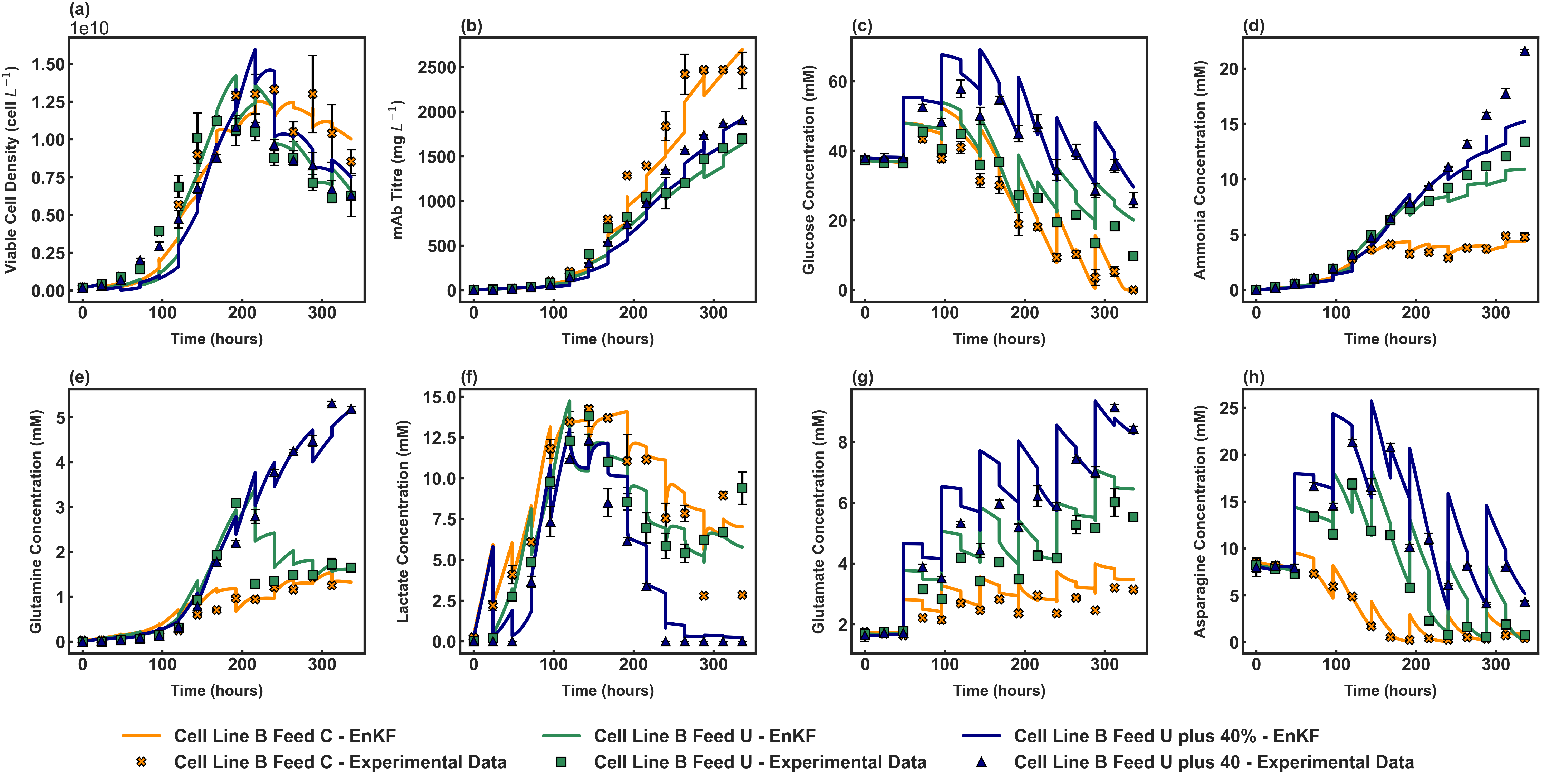
EnKF-based state estimation model transfer across cell line and feeding strategies for Cell Line B cultures.

##### Effect of excess feeding on biomass yield and nitrogen metabolism

To assess how elevated glucose and asparagine in the in-house feeds influence growth in the new CHO-GS cell line, the temporal evolution of biomass syields from glucose *Y*_*X*,*Glc*_ and asparagine *Y*_*X*,*Asn*_ was examined. As shown in Figure 6 (e), *Y*_*X*,*Glc*_ exhibits broad ensemble spread for all conditions, with only modest changes in the parameter mean over time. This indicates that *Y*_*X*,*Glc*_ is relatively insensitive to variations in glucose availability and does not strongly influence growth predictions.

**Figure 6.**
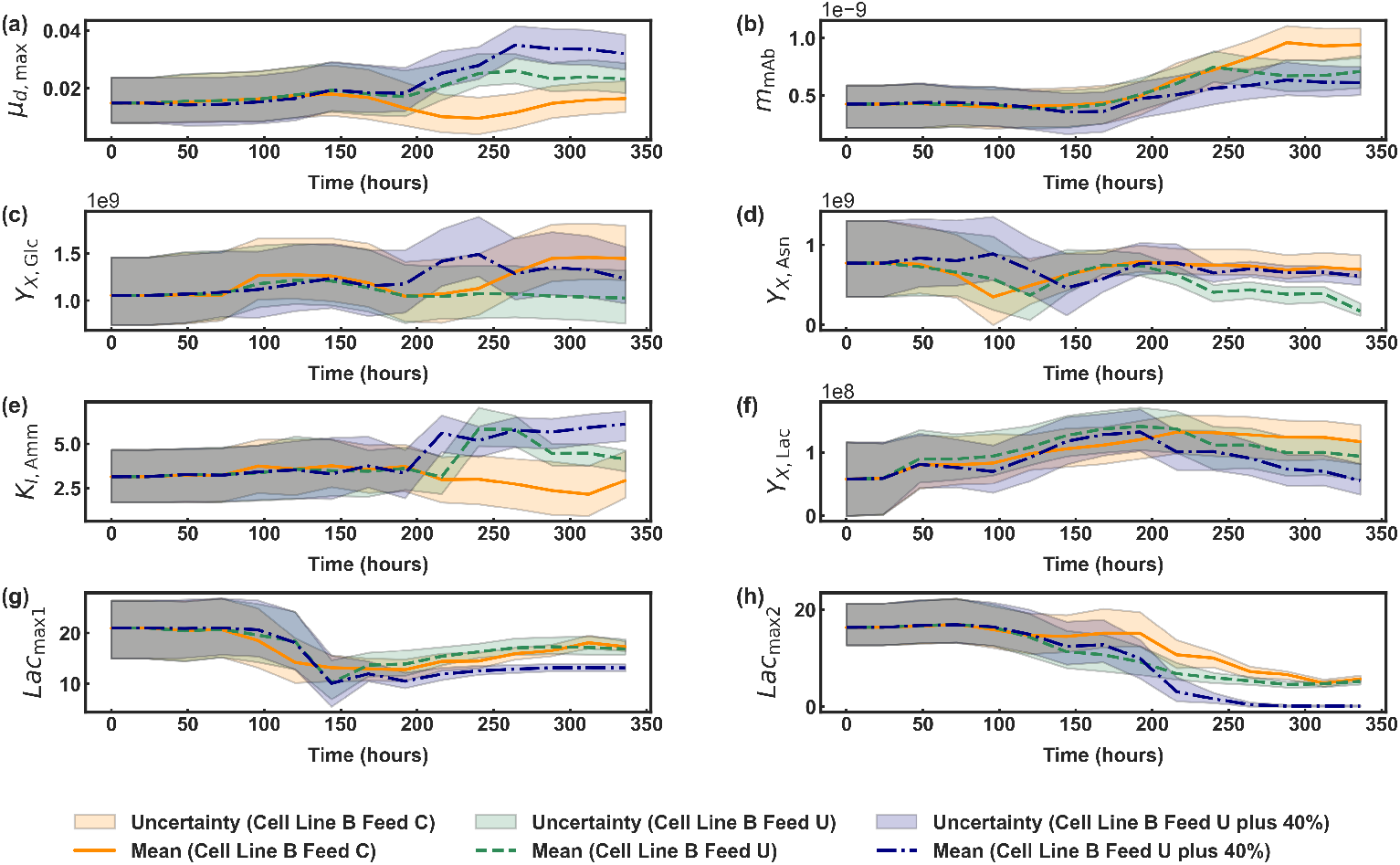
Time evolution of estimated model parameters across cell line and feeding strategies for Cell Line B cultures.

In contrast, the *Y*_*X*,*Asn*_ ensemble spread reduces rapidly, indicating that the filter gains confidence quickly about the contribution of asparagine to biomass formation. *Y*_*X*,*Asn*_ is initially highest in Feed U plus 40%, reflecting its elevated concentration. Over time, this parameter declines in Feed U with reduced uncertainty, indicating growing model confidence that asparagine contributes less to biomass. Interestingly, *Y*_*X*,*Asn*_ in Feed U plus 40% drops below that of Feed C by the end, despite significantly higher asparagine availability and uptake rate. This disparity suggests rerouting of excess asparagine through alternative nitrogen pathways. Previous studies have proposed that, under nitrogen-rich conditions, asparagine synthetase may catalyse a reaction analogous to asparaginase, converting asparagine and glutamate into aspartate and glutamine [63]. While not conventionally reported in mammalian systems, this reaction has been included in CHO metabolic reconstructions to account for elevated ammonia accumulation under nitrogen-rich conditions [58, 64]. The parameter trajectory of *Y*_*X*,*Asn*_ thus offers indirect evidence of such metabolic shifts, revealing insight not captured by concentration data alone.

To further investigate the impact of excess asparagine feeding, the ammonia inhibition parameter *K*_*I*,*Amm*_ was evaluated. In Feed C, this parameter remains near its initial value with broad uncertainty, consistent with its use in model calibration. In contrast, both in-house feeds exhibit a sharp increase in *K*_*I*,*Amm*_ entering stationary phase, accompanied by a rapid drop in uncertainty. This trend aligns with the timing of ammonia accumulation and declining *Y*_*X*,*Asn*_, indicating that excess nitrogen progressively limits growth in later stages.

##### Understanding lactate metabolism beyond concentration profiles

To understand how lactate metabolism differs across feeding strategies, the biomass yield on lactate *Y*_*X*,*Lac*_ and two kinetic parameters *Lac*_*max*1_ and *Lac*_*max*2_ were analysed. While concentration profiles confirm net lactate consumption under all conditions, they cannot distinguish whether this uptake supports biomass growth or other cellular functions.The parameter trajectories provide a clearer picture. All three conditions show elevated *Y*_*X*,*Lac*_ early in culture, suggesting enhanced growth-coupled lactate utilisation in Cell Line B. However, only Feed C maintains high yields in later stages, while both in-house formulations show a decline. This supports the hypothesis that, lactate may be increasingly rerouted toward non-growth-associated functions such as redox balancing under nutrient-rich conditions [65].

This interpretation is further supported by the trajectories of activation constants for lactate consumption *Lac*_*max*1_ and *Lac*_*max*2_. *Lac*_*max*2_ declines across all feeds but most sharply under Feed U plus 40%, in line with complete lactate depletion. In the same condition, *Lac*_*max*1_ also drops steadily, indicating sustained consumption. In contrast, under Feed C and Feed U, *Lac*_*max*1_ rises again, suggesting some lactate accumulation remains despite overall uptake. Additionally, although the concentration profiles indicate net lactate accumulation throughout the early phase, the parameter trajectories indicate an underlying metabolic shift. Both *Lac*_max1_ and *Lac*_max2_ begin to decline as early as day 4, well before glucose or amino acid depletion. This behaviour mirrors previous observations in Cell Line B batch cultures [66] and supports literature findings that lactate consumption can also be triggered by high extracellular lactate rather than nutrient limitation [67, 68].

### 3.3 Long Term Prediction Transferring Cell-line and Feeding Strategy

In addition to recursive estimation, the EnKF can also be applied for long-term prediction. As an illustrative example, the parameter ensemble is updated every time a measurement becomes available and used to simulate culture trajectories from the respective time point until harvest. These simulations are performed open-loop, meaning no further measurement updates are incorporated beyond the current step. At each time point, the forecasts therefore represent the best available process knowledge given the parameter values inferred up to that stage. As additional data become available, parameter estimates are updated leading to progressively improved forecasts. Figure 7 illustrates the long-term prediction performance of EnKF when applied to Cell Line B cultured with Feed U plus 40%, a system that differs significantly from the Cell Line A baseline both in cell line and feeding strategy. By day 7, EnKF predictions already substantially improved upon the baseline mechanistic model for most states, including glucose, glutamine, lactate, and glutamate. By day 9, accurate forecasts are achieved for glucose, glutamine, glutamate, and asparagine. By day 12, all state trajectories align closely with those obtained using the full dataset, confirming the framework’s capacity to refine harvest-day outcomes in real time. The corresponding long-term prediction results for the remaining datasets are provided in Supplementary Information F. Such progressively improving forecasts can support model-predictive control, where the updated parameter ensemble informs the optimisation of future control actions using the most recent process knowledge.

**Figure 7.**
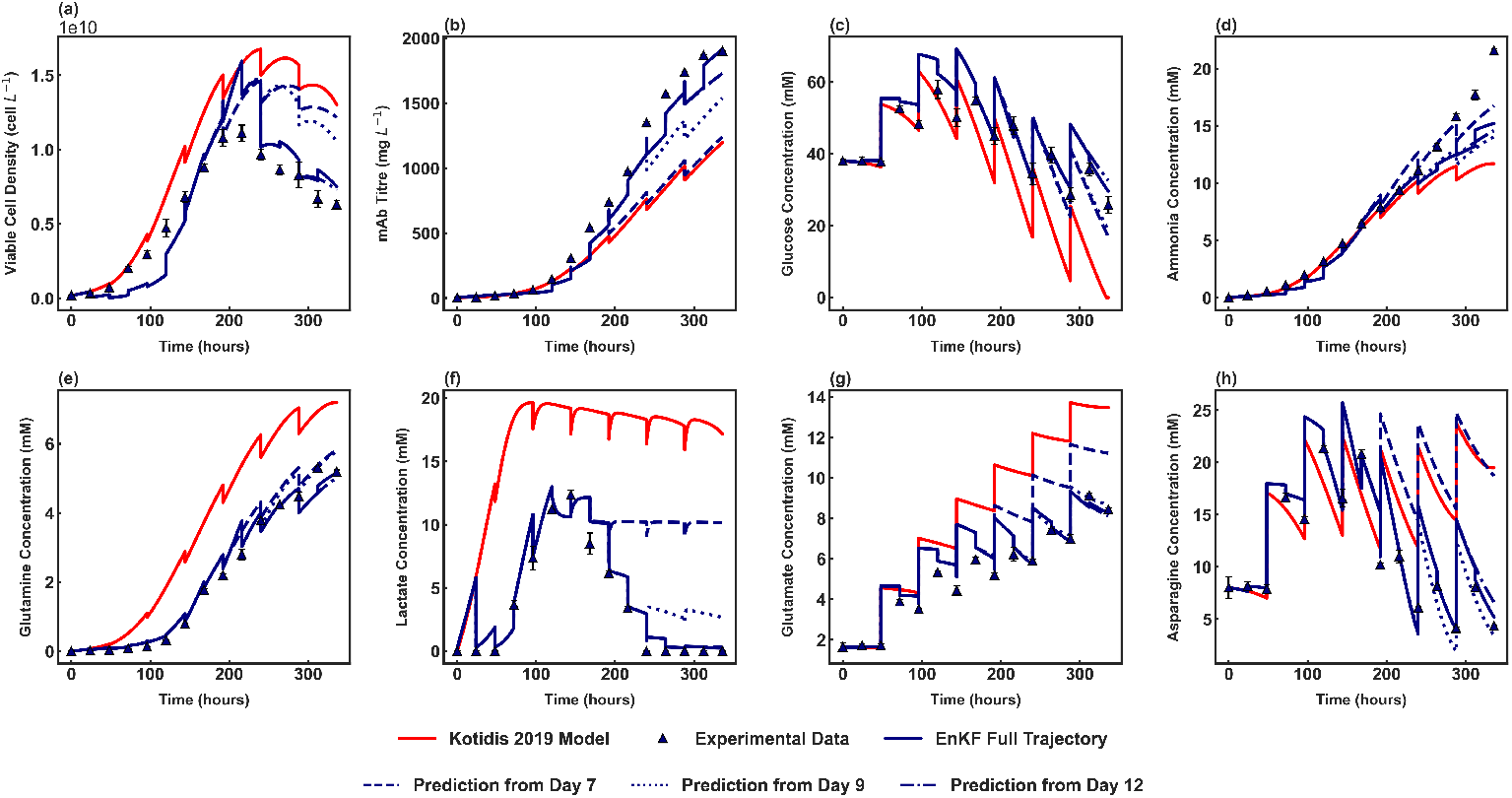
Long term EnKF predictions of Cell Line B Feed U plus 40% initiated from day 7 (dash), day 9 (dotted), and day 12 (dashed-dot).

However, the timing at which accurate long-term prediction becomes possible is case-dependent. It reflects both the degree of deviation from the base model and the availability of information on key metabolic shifts. For instance, the sharp decline in viable cell density near the end of culture is not reflected in predictions from day 7 or day 9, as this behaviour had not yet emerged in the available data. Since the EnKF builds upon a mechanistic model originally developed for a different system, its ability to adapt relies on the progressive assimilation of system-specific measurements. Thus, while the framework is powerful in adapting to new conditions, its predictive accuracy is fundamentally determined by the timing and availability of incoming data.

## 4 Conclusions

This work establishes the EnKF as a dynamic and mechanistically grounded framework for knowledge transfer in upstream bioprocess modelling, allowing a system-specific mechanistic model to be progressively adapted to new conditions without reparametrisation. A central contribution lies in the dynamic evolution of parameter ensembles, which provide a temporally resolved view of how the model adapts to the new process. Unlike traditional approaches that treat model parameters as static, the EnKF framework tracks how parameter estimates evolve in response to real-time measurements. This time-resolved sensitivity analysis reveals which parameters are the most influential under new conditions, and when their impact on model behaviour becomes significant. Importantly, this is achieved without introducing artificial variability through parameter covariance inflation, thereby ensuring that changes in ensemble spread arise from real process data only.

Preserving mechanistic interpretability is particularly valuable for knowledge transfer, as it allows models to explain system behaviour beyond predictive performance. While hybrid and black-box approaches often prioritise generalisability, recent studies demonstrate that full mechanistic transparency can still be retained across diverse transfer contexts. Richelle *et al*. demonstrated transfer from fed-batch to perfusion by simplifying model structure and introducing a ‘catch-all’ by-product variable to accommodate changes in operating mode [69]. Arndt *et al*. developed a scale-up workflow that quantified and compared model parameter distributions across bioreactor volumes, revealing when transfer is valid and when scale-dependent reparametrisation is necessary [70]. More recently, Lu *et al*. incorporated time-series transcriptomics into a multi-cell-line learning (MCL) framework providing enzyme-level metabolic insight across CHO clones, though requiring high-performance computing and rich omics datasets [22]. This work complements these efforts by introducing time-resolved parameter trajectories, with the EnKF treating biologically meaningful kinetic parameters as dynamic quantities that evolve in response to process measurements. Compared with MCL, the presented method offers a computationally efficient alternative using only routine extracellular measurements, particularly suited for early process development where omics data may not be available.

While this method offers notable advantages in flexibility and interpretability, it relies on two key assumptions. First, the method is designed to capture process-specific differences through parameter adaptation, without changing the underlying model structure. Second, the evolution of parameters is inherently conditioned on the initial ensemble drawn from the base system. In highly nonlinear systems, multiple parameter combinations may yield similar outputs, and the estimated trajectory represents one of the plausible solutions. These assumptions do not undermine the utility of the method but are essential to consider when interpreting biological meaning. Provided the model structure remains appropriate, the dynamic adaptation of parameters still offers valuable insight into system behaviour and sensitivity, especially when full reparametrisation is not feasible.

As the first demonstration EnKF dual state and parameter estimation in bioprocessing literatures, this work shows how dynamic model adaptation based on routine extracellular measurements can reveal system-specific behaviour without requiring full reparametrisation. The resulting dynamic parameter trajectories not only increase the flexibility of transferring mechanistic models across systems, but also provide biologically meaningful insight into process adaptation that extends beyond what is observable from concentration profiles alone. Beyond its methodological contributions, this framework offers a practical and interpretable route for reusing mechanistic models in new bioprocess settings where data are limited and timelines are constrained. From a regulatory perspective, the preservation of mechanistic structure and the transparent evolution of model parameters can support scientific justification for process comparability and risk assessment during tech transfer. In this context, the EnKF establishes itself as a robust and accessible tool for supporting knowledge-driven decision-making across development stages in upstream bioprocessing.

## 5 Acknowledgments

Three authors have no competing interest to declare. Author contributions are as follows. LY: Conceptualization, Methodology, Formal Analysis, Writing - Original Draft; ARC: Conceptualization, Supervision, Writing - Review and Editing; CK: Conceptualization, Supervision, Writing - Review and Editing.

## A Mechanistic Model from Kotidis *et al*. 2019

### CHO cell growth and death

The CHO cell growth and death kinetics were represented using Monod-type relationships, where cell proliferation depends on essential nutrients and is inhibited by metabolic by-products. The dynamic balances describing the culture volume and viable cell population are given by:

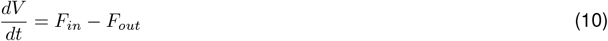

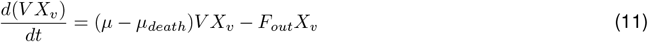

where *V* (L) denotes the culture volume, *t* (h) the cultivation time, and *F*_*in*_ and *F*_*out*_ (L·h^−1^) the feeding and sampling flow rates. *X*_*v*_ (cell·L^−1^) is the viable cell density, while *µ* and *µ*_*death*_ (h^−1^) denote the specific growth and death rates.

Extracellular concentration (mM) of glucose [*Glc*] and asparagine [*Asn*] were modelled as the primary limiting substrates supporting cell proliferation. In the original Kotidis *et al*. (2019) model, uridine [*Urd*] acted as an inhibitory and cytotoxic factor influencing both *µ* and *µ*_death_. Since the current datasets did not involve uridine supplementation, these dependencies were excluded, and inhibitory effects were represented only through lactate [*Lac*] and ammonia [*Amm*] accumulation. The specific growth rate was expressed as:

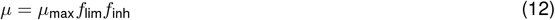

where *µ*_max_ (h^−1^) is the maximum specific growth rate, *f*_lim_ represents nutrient limitation, and *f*_inh_ represents inhibition by metabolic by-products:

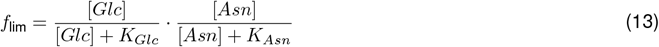

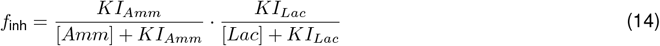

Here, *K*_*Glc*_ and *K*_*Asn*_ (mM) denote the Monod constants for glucose and asparagine uptake, while *KI*_*Amm*_ and *KI*_*Lac*_ (mM) are inhibition constants for ammonia and lactate. The cell death rate was modelled as a function of ammonia toxicity:

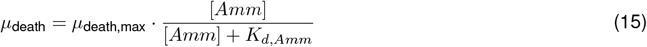

where *µ*_death,max_ (h^−1^) is the maximum specific death rate and *K*_*d*,*Amm*_ (mM) the ammonia toxicity constant.

**Table S1:** Fitted parameters for the CHO cell growth and death model.

| Parameter | Description | Value | Unit |
| --- | --- | --- | --- |
| $\mu_{max}$ | Maximum specific growth rate | 0.065 | h<sup>-1</sup> |
| $\mu_{death,max}$ | Maximum specific cell death rate | 0.015 | h<sup>-1</sup> |
| $K_{Glc}$ | Monod constant for glucose uptake | 14.04 | mM |
| $K_{Asn}$ | Monod constant for asparagine uptake | 2.62 | mM |
| $K I_{Amm}$ | Inhibition constant for ammonia | 3.17 | mM |
| $K I_{Lac}$ | Inhibition constant for lactate | 1000 | mM |
| $K_{d,Amm}$ | Death constant for ammonia toxicity | 14.28 | mM |

### CHO cell metabolism

The metabolic framework describes extracellular metabolite and amino acid dynamics through material balance equations. Each balance accounts for feeding, sampling, and cell-specific production or consumption rates as follows:

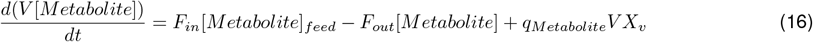

where [*Metabolite*] (mM) represents the extracellular concentration, [*Metabolite*]_*feed*_ (mM) the feed concentration, and *q*_*Metabolite*_ (mmol·cell^−1^·h^−1^) the specific production or consumption rate. The following subsections summarise the kinetic expressions used for each metabolite.

#### Glucose

In the original Kotidis *et al*. 2019 model, glucose uptake included a galactose-dependent competitive term to capture flux diversion between the two sugars. As no galactose supplementation was present in the datasets used in this study, this term was omitted, and glucose uptake was modelled solely as a function of growth and maintenance demands:

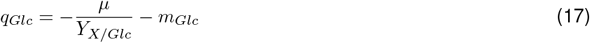

where *Y*_*X/Glc*_ (cell·mmol^−1^) is the yield of biomass on glucose and *m*_*Glc*_ (mmol·cell^−1^·h^−1^) is the maintenance coefficient for glucose.

#### Glutamine

The specific glutamine rate was expressed as:

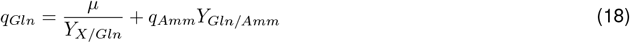

where *Y*_*X/Gln*_ (cell·mmol^−1^) is the yield of biomass on glutamine and *Y*_*Gln/Amm*_ (mmol·mmol^−1^) is the yield of glutamine from ammonia.

#### Lactate

Lactate dynamics incorporated both production and consumption behaviour:

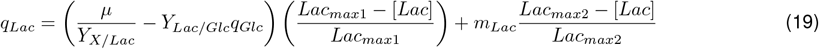

where *Y*_*X/Lac*_ (cell·mmol^−1^) is the yield of biomass on lactate, *Y*_*Lac/Glc*_ (mmol·mmol^−1^) is the yield of lactate from glucose consumption towards the glycolysis pathway and through the synthesis of pyruvate, *Lac*_*max*1_ and *Lac*_*max*2_ (mM) are the activation constants for lactate consumption, and *m*_*Lac*_ (mmol·cell^−1^·h^−1^) is the maintenance coefficient for the participation of lactate in other metabolic pathways of the cell.

#### Ammonia

Ammonia accumulation was described as:

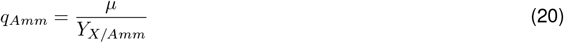

where *Y*_*X/Amm*_ (cell·mmol^−1^) is the yield of biomass on ammonia.

#### Glutamate

Glutamate consumption was expressed as:

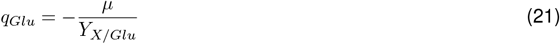

where *Y*_*X/Glu*_ (cell·mmol^−1^) is the yield of biomass on glutamate.

#### Asparagine and Aspartate

Asparagine uptake and its coupling with aspartate synthesis were modelled as:

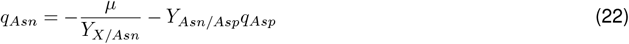

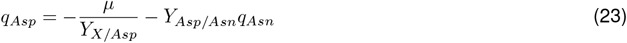

where *Y*_*X/Asn*_ and *Y*_*X/Asp*_ (cell·mmol^−1^) are the yields of biomass on asparagine and aspartate, respectively, while *Y*_*Asn/Asp*_ and *Y*_*Asp/Asn*_ (mmol·mmol^−1^) are the interconversion yields between asparagine and aspartate.

#### Monoclonal antibody synthesis

Antibody production was described by a material balance on the extracellular antibody concentration:

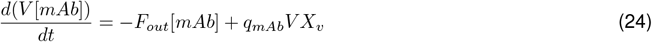

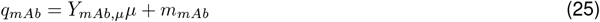

where *Y*_*mAb*,*µ*_ (mg·cell^−1^) is the yield of monoclonal antibody on cell growth and *m*_*mAb*_ (mg·cell^−1^·h^−1^) is the non-growth associated mAb production rate.

**Table S2:** Fitted parameters for the CHO cell metabolism and monoclonal antibody synthesis models.

| Parameter | Description | Value | Unit |
| --- | --- | --- | --- |
| $Y_{X/Glc}$ | Yield of biomass on glucose | $1.01 \times 10^9$ | $\text{cell}\cdot\text{mmol}^{-1}$ |
| $m_{Glc}$ | Maintenance coefficient for glucose | $3.43 \times 10^{-11}$ | $\text{mmol}\cdot\text{cell}^{-1}\cdot\text{h}^{-1}$ |
| $Y_{X/Gln}$ | Yield of biomass on glutamine | $4.64 \times 10^9$ | $\text{cell}\cdot\text{mmol}^{-1}$ |
| $Y_{Gln/Amm}$ | Yield of glutamine from ammonia | 0.10 | $\text{mmol}\cdot\text{mmol}^{-1}$ |
| $Y_{X/Lac}$ | Yield of biomass on lactate | $5.46 \times 10^7$ | $\text{cell}\cdot\text{mmol}^{-1}$ |
| $Y_{Lac/Glc}$ | Yield of lactate from glucose | 1.56 | $\text{mmol}\cdot\text{mmol}^{-1}$ |
| $Lac_{max1}$ | Activation constant for lactate consumption | 21.20 | mM |
| $Lac_{max2}$ | Activation constant for lactate consumption | 16.00 | mM |
| $m_{Lac}$ | Maintenance coefficient for lactate | $1.87 \times 10^{-10}$ | $\text{mmol}\cdot\text{cell}^{-1}\cdot\text{h}^{-1}$ |
| $Y_{X/Amm}$ | Yield of biomass on ammonia | $2.36 \times 10^9$ | $\text{cell}\cdot\text{mmol}^{-1}$ |
| $Y_{X/Glu}$ | Yield of biomass on glutamate | $1.46 \times 10^{10}$ | $\text{cell}\cdot\text{mmol}^{-1}$ |
| $Y_{X/Asn}$ | Yield of biomass on asparagine | $7.68 \times 10^8$ | $\text{cell}\cdot\text{mmol}^{-1}$ |
| $Y_{Asn/Asp}$ | Yield of asparagine from aspartate | 0.10 | $\text{mmol}\cdot\text{mmol}^{-1}$ |
| $Y_{X/Asp}$ | Yield of biomass on aspartate | $3.59 \times 10^9$ | $\text{cell}\cdot\text{mmol}^{-1}$ |
| $Y_{Asp/Asn}$ | Yield of aspartate from asparagine | 0.13 | $\text{mmol}\cdot\text{mmol}^{-1}$ |
| $Y_{mAb,\mu}$ | Yield of monoclonal antibody on cell growth | 3.39 | $\text{pg}\cdot\text{cell}^{-1}$ |
| $m_{mAb}$ | Non-growth associated mAb production rate | $4.10 \times 10^{-1}$ | $\text{pg}\cdot\text{cell}^{-1}\cdot\text{h}^{-1}$ |

## B Measurement and Process Noises

### Measurement Noise Tuning

A fixed scalar inflation factor of *k*_*R*_ = 1 was applied across all conditions to scale the measurement noise.

**Table S3:** Measurement covariance (*R*) values defining the observation noise variance for each dataset.

| State | Cell Line A<br>Shake Flask | Cell Line A<br>Bioreactor<br>36.5°C | Cell Line A<br>Bioreactor<br>32°C | Cell Line B<br>Feed C | Cell Line B<br>Feed U | Cell Line B<br>Feed U + 40% |
| --- | --- | --- | --- | --- | --- | --- |
| Xv | $3.0 \times 10^{17}$ | $5.0 \times 10^{17}$ | $3.0 \times 10^{17}$ | $3.0 \times 10^{17}$ | $3.0 \times 10^{17}$ | $3.0 \times 10^{17}$ |
| mAb | $1.5 \times 10^3$ | $1.5 \times 10^3$ | $1.5 \times 10^3$ | $1.5 \times 10^3$ | $1.5 \times 10^3$ | $1.5 \times 10^3$ |
| Glc | 5.0 | 5.0 | 5.0 | 5.0 | 5.0 | 5.0 |
| Amm | 0.1 | 0.1 | 0.2 | 0.1 | 0.1 | 0.2 |
| Gln | 0.1 | 0.1 | 0.1 | 0.1 | 0.1 | 0.1 |
| Lac | 1.2 | 1.2 | 0.5 | 0.2 | 0.2 | 0.2 |
| Glu | 0.004 | 0.004 | 0.04 | 0.08 | 0.08 | 0.04 |
| Asn | 0.25 | 0.25 | 0.25 | 0.025 | 0.025 | 0.025 |

### Process Noise Tuning

A fixed scalar inflation factor of *k*_*Q*_ = 1 *×* 10 ^−9^ was applied across all conditions to scale the process noise.

**Table S4:** Process covariance (*Q*) values defining the process noise variance for each dataset.

| State | Cell Line A<br>Shake Flask | Cell Line A<br>Bioreactor<br>36.5°C | Cell Line A<br>Bioreactor<br>32°C | Cell Line B<br>Feed C | Cell Line B<br>Feed U | Cell Line B<br>Feed U + 40% |
| --- | --- | --- | --- | --- | --- | --- |
| Xv | $1.0 \times 10^{18}$ | $1.0 \times 10^{19}$ | $2.0 \times 10^{19}$ | $2.0 \times 10^{19}$ | $2.0 \times 10^{19}$ | $2.0 \times 10^{19}$ |
| mAb | $1.0 \times 10^4$ | $1.0 \times 10^5$ | $1.0 \times 10^5$ | $1.0 \times 10^5$ | $1.0 \times 10^5$ | $1.0 \times 10^5$ |
| Glc | $1.6 \times 10^1$ | $1.6 \times 10^1$ | $9.0 \times 10^2$ | $9.6 \times 10^1$ | $9.6 \times 10^1$ | $9.6 \times 10^1$ |
| Amm | 1.0 | 1.0 | 6.0 | 6.0 | 6.0 | 10.0 |
| Gln | 2.0 | 4.0 | 4.0 | 4.0 | 4.0 | 4.0 |
| Lac | 1.2 | 1.2 | 5.0 | 5.0 | 5.0 | 5.0 |
| Glu | 0.2 | 0.2 | 1.6 | 1.6 | 1.6 | 1.4 |
| Asn | 0.8 | 0.8 | 5.0 | 3.0 | 3.0 | 3.0 |

### C Parameter Covariances

**Table S5:** Mean parameter values and ensemble covariances used for EnKF initialization. Initial parameter mean obtained from Kotidis *et al*. 2019 model parameters [56].

| Parameter | Initial Parameter Mean | Unit | Parameter Covariance |
| --- | --- | --- | --- |
| $\mu_{max}$ | 0.065 | $\text{h}^{-1}$ | 0.02 |
| $\mu_{d,max}$ | 0.02 | $\text{h}^{-1}$ | 0.0035 |
| $K_{Glc}$ | 14.04 | mM | 4.20 |
| $K_{Asn}$ | 2.62 | mM | 0.50 |
| $KI_{Lac}$ | 1000 | mM | 150 |
| $KI_{Amm}$ | 3.17 | mM | 0.69 |
| $K_{d,Amm}$ | 14.28 | mM | 2.50 |
| $m_{Lac}$ | $1.87 \times 10^{-10}$ | $\text{mmol} \cdot \text{cell}^{-1} \cdot \text{h}^{-1}$ | $2.00 \times 10^{-11}$ |
| $m_{Glc}$ | $3.43 \times 10^{-11}$ | $\text{mmol} \cdot \text{cell}^{-1} \cdot \text{h}^{-1}$ | $3.10 \times 10^{-12}$ |
| $Lac_{max1}$ | 21.20 | mM | 3.00 |
| $Lac_{max2}$ | 16.00 | mM | 2.00 |
| $Y_{X,Glc}$ | $1.01 \times 10^9$ | $\text{cell} \cdot \text{mmol}^{-1}$ | $1.77 \times 10^8$ |
| $Y_{X,Gln}$ | $4.64 \times 10^9$ | $\text{cell} \cdot \text{mmol}^{-1}$ | $6.03 \times 10^8$ |
| $Y_{X,Glu}$ | $1.46 \times 10^{10}$ | $\text{cell} \cdot \text{mmol}^{-1}$ | $3.30 \times 10^9$ |
| $Y_{X,Lac}$ | $5.46 \times 10^7$ | $\text{cell} \cdot \text{mmol}^{-1}$ | $2.57 \times 10^7$ |
| $Y_{X,Amm}$ | $2.36 \times 10^9$ | $\text{cell} \cdot \text{mmol}^{-1}$ | $6.02 \times 10^8$ |
| $Y_{X,Asn}$ | $7.68 \times 10^8$ | $\text{cell} \cdot \text{mmol}^{-1}$ | $2.08 \times 10^8$ |
| $Y_{X,Asp}$ | $3.59 \times 10^9$ | $\text{cell} \cdot \text{mmol}^{-1}$ | $8.07 \times 10^8$ |
| $Y_{Gln,Amm}$ | 0.10 | $\text{mmol} \cdot \text{mmol}^{-1}$ | 0.03 |
| $Y_{Lac,Glc}$ | 1.56 | $\text{mmol} \cdot \text{mmol}^{-1}$ | 0.15 |
| $Y_{Asp,Asn}$ | 0.13 | $\text{mmol} \cdot \text{mmol}^{-1}$ | 0.01 |
| $Y_{Asn,Asp}$ | 0.10 | $\text{mmol} \cdot \text{mmol}^{-1}$ | 0.01 |
| $m_{mAb}$ | $4.10 \times 10^{-10}$ | $\text{mg} \cdot \text{cell}^{-1} \cdot \text{h}^{-1}$ | $9.05 \times 10^{-11}$ |
| $Y_{mAb,\mu}$ | $3.39 \times 10^{-9}$ | $\text{mg} \cdot \text{cell}^{-1}$ | $3.10 \times 10^{-10}$ |

## D Parameter Sensitivity Analysis

**Table S6:** Parameter uncertainty reduction based on ensemble spread, with top 6 most sensitive parameters highlighted for each dataset.

| Parameter | Cell Line A<br>Shake Flask<br>(%) | Cell Line A<br>Bioreactor<br>36.5°C (%) | Cell Line A<br>Bioreactor<br>32°C (%) | Cell Line B<br>Feed C (%) | Cell Line B<br>Feed U (%) | Cell Line B<br>Feed U + 40%<br>(%) |
| --- | --- | --- | --- | --- | --- | --- |
| $\mu_{\max}$ | 66.9 | 54.3 | 61.6 | 45.1 | 80.3 | 65.2 |
| $\mu_{d,\max}$ | 31.5 | 36.2 | 25.5 | 22.4 | 34.4 | 31.1 |
| $K_{\text{Glc}}$ | 37.0 | 35.3 | 29.4 | 28.9 | 29.6 | 32.1 |
| $K_{\text{Asn}}$ | 35.3 | 34.7 | 25.8 | 31.9 | 31.3 | 33.3 |
| $K I_{\text{Lac}}$ | 32.0 | 41.1 | 35.4 | 27.0 | 26.6 | 31.2 |
| $K I_{\text{Amm}}$ | 34.9 | 43.1 | 37.9 | 29.0 | 69.4 | 52.7 |
| $K_{d,\text{Amm}}$ | 30.0 | 29.3 | 33.6 | 25.6 | 29.2 | 42.4 |
| $m_{\text{Lac}}$ | 27.4 | 25.6 | 25.4 | 22.6 | 30.8 | 38.8 |
| $m_{\text{Glc}}$ | 41.4 | 62.3 | 25.7 | 38.0 | 37.3 | 40.3 |
| $Lac_{\max 1}$ | 59.7 | 58.5 | 64.2 | 85.2 | 82.0 | 87.0 |
| $Lac_{\max 2}$ | 48.6 | 63.4 | 86.6 | 85.8 | 88.8 | 96.1 |
| $Y_{X,\text{Glc}}$ | 43.8 | 47.3 | 35.5 | 28.3 | 29.9 | 30.8 |
| $Y_{X,\text{Gln}}$ | 30.3 | 29.9 | 36.8 | 19.2 | 23.1 | 27.3 |
| $Y_{X,\text{Glu}}$ | 41.1 | 81.4 | 57.8 | 56.8 | 62.9 | 80.0 |
| $Y_{X,\text{Lac}}$ | 64.6 | 60.3 | 59.2 | 54.4 | 58.4 | 62.8 |
| $Y_{X,\text{Amm}}$ | 68.9 | 71.0 | 35.9 | 41.4 | 63.7 | 57.8 |
| $Y_{X,\text{Asn}}$ | 74.3 | 67.2 | 43.5 | 74.2 | 85.0 | 83.7 |
| $Y_{X,\text{Asp}}$ | 8.8 | 11.4 | 9.4 | 15.7 | 16.7 | 21.1 |
| $Y_{\text{Gln},\text{Amm}}$ | 20.2 | 26.0 | 24.6 | 25.3 | 27.3 | 29.7 |
| $Y_{\text{Lac},\text{Glc}}$ | 21.3 | 26.5 | 29.8 | 22.9 | 23.7 | 21.7 |
| $Y_{\text{Asp},\text{Asn}}$ | 22.1 | 34.6 | 33.6 | 21.3 | 22.7 | 27.0 |
| $Y_{\text{Asn},\text{Asp}}$ | 9.5 | 26.5 | 24.5 | 24.3 | 25.6 | 25.1 |
| $m_{\text{mAb}}$ | 43.8 | 37.0 | 33.0 | 24.2 | 29.5 | 34.3 |
| $Y_{\text{mAb},\mu}$ | 13.4 | 28.5 | 31.3 | 30.6 | 28.4 | 27.6 |

## E RMSE of EnKF Compared with Experimental Data

**Table S7:** Average root mean squared error (RMSE) between EnKF-predicted and experimental (benchmark) state trajectories across datasets.

| State | Cell Line A<br>Shake Flask<br>(%) | Cell Line A<br>Bioreactor<br>36.5°C (%) | Cell Line A<br>Bioreactor<br>32°C (%) | Cell Line B<br>Feed C (%) | Cell Line B<br>Feed U (%) | Cell Line B<br>Feed U + 40%<br>(%) |
| --- | --- | --- | --- | --- | --- | --- |
| Xv | $3.14 \times 10^8$ | $1.63 \times 10^9$ | $2.08 \times 10^9$ | $1.55 \times 10^9$ | $1.64 \times 10^9$ | $1.84 \times 10^9$ |
| mAb | 9.89 | $1.03 \times 10^2$ | $9.31 \times 10^1$ | $2.05 \times 10^2$ | $1.00 \times 10^2$ | $1.33 \times 10^2$ |
| Glc | 1.11 | 7.11 | 8.63 | 3.25 | 4.32 | 3.34 |
| Amm | 0.33 | 0.86 | 0.62 | 0.49 | 0.92 | 1.60 |
| Gln | 0.16 | 0.70 | 1.08 | 0.24 | 0.38 | 0.27 |
| Lac | 1.44 | 2.28 | 3.62 | 2.21 | 2.05 | 2.24 |
| Glu | 0.15 | 0.16 | 0.46 | 0.39 | 0.57 | 0.60 |
| Asn | 0.19 | 0.26 | 0.73 | 0.48 | 1.32 | 1.58 |

## F Long Term Prediction

**Figure S1:**
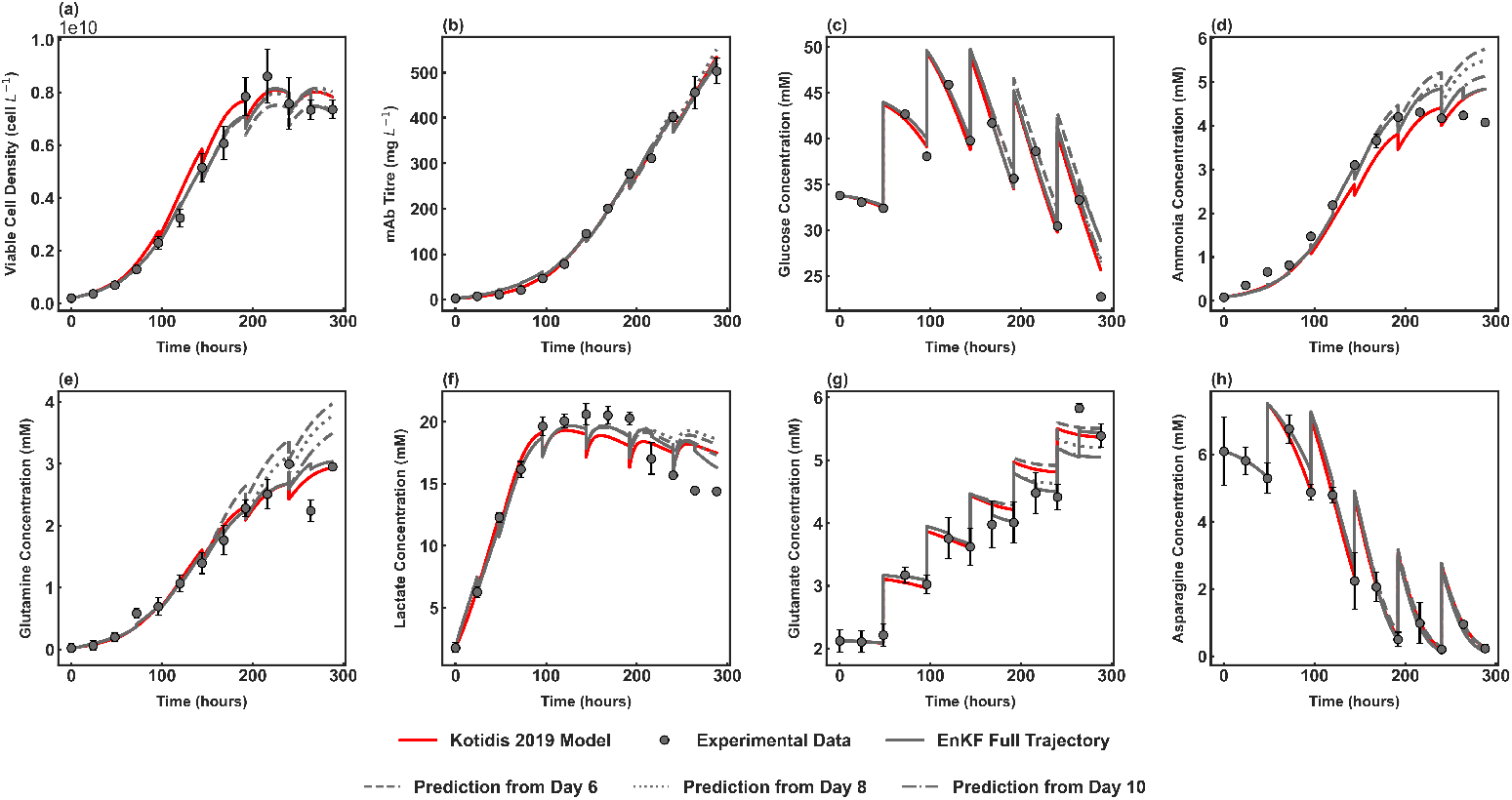
Long term EnKF predictions of Cell Line A Shake Flask at 36.5 °C initiated from Day 7 (dash), Day 9 (dotted), and Day 12 (dashed-dot).

**Figure S2:**
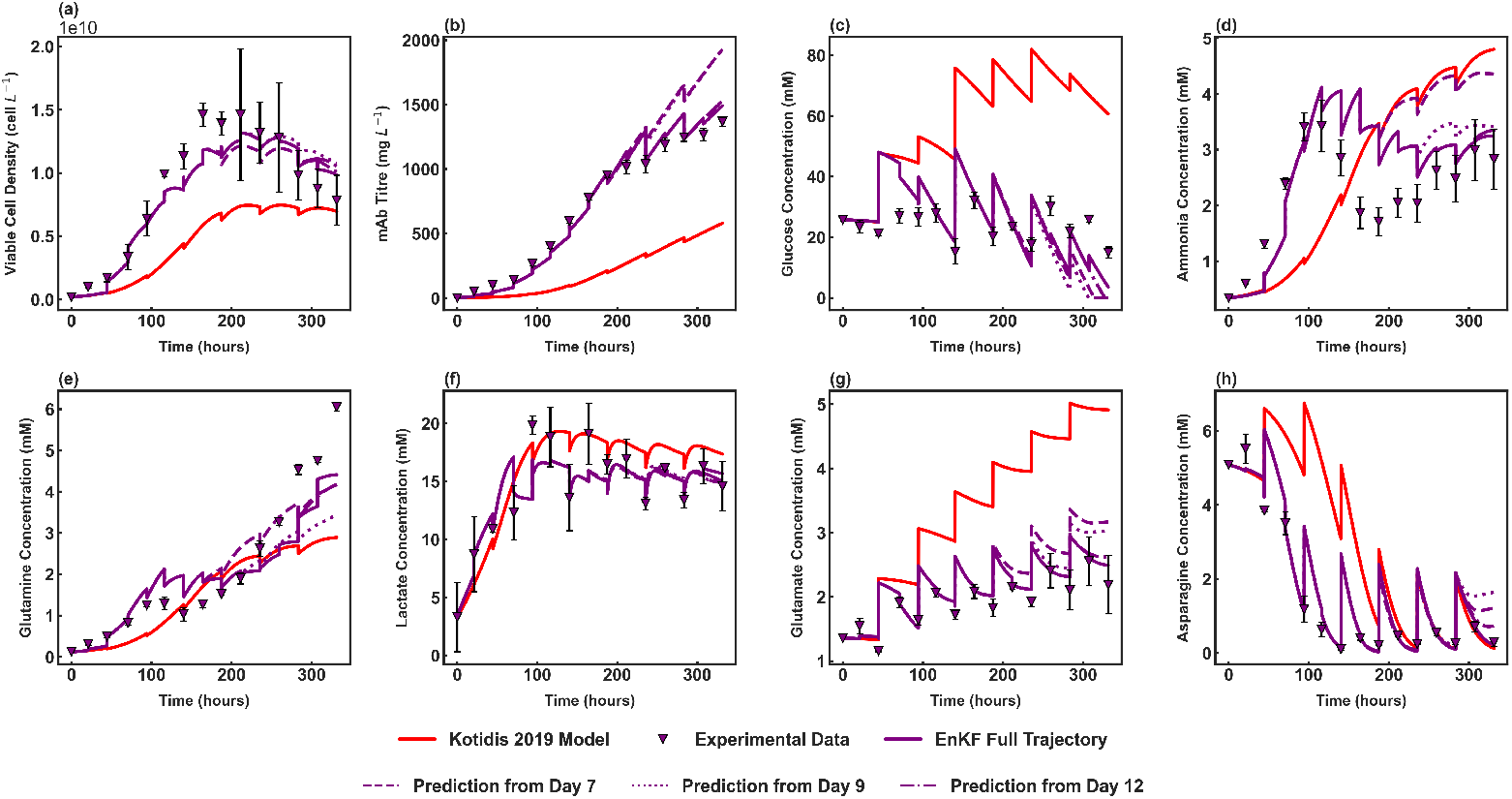
Long term EnKF predictions of Cell Line A Bioreactor at 36.5 °C initiated from Day 7 (dash), Day 9 (dotted), and Day 12 (dashed-dot).

**Figure S3:**
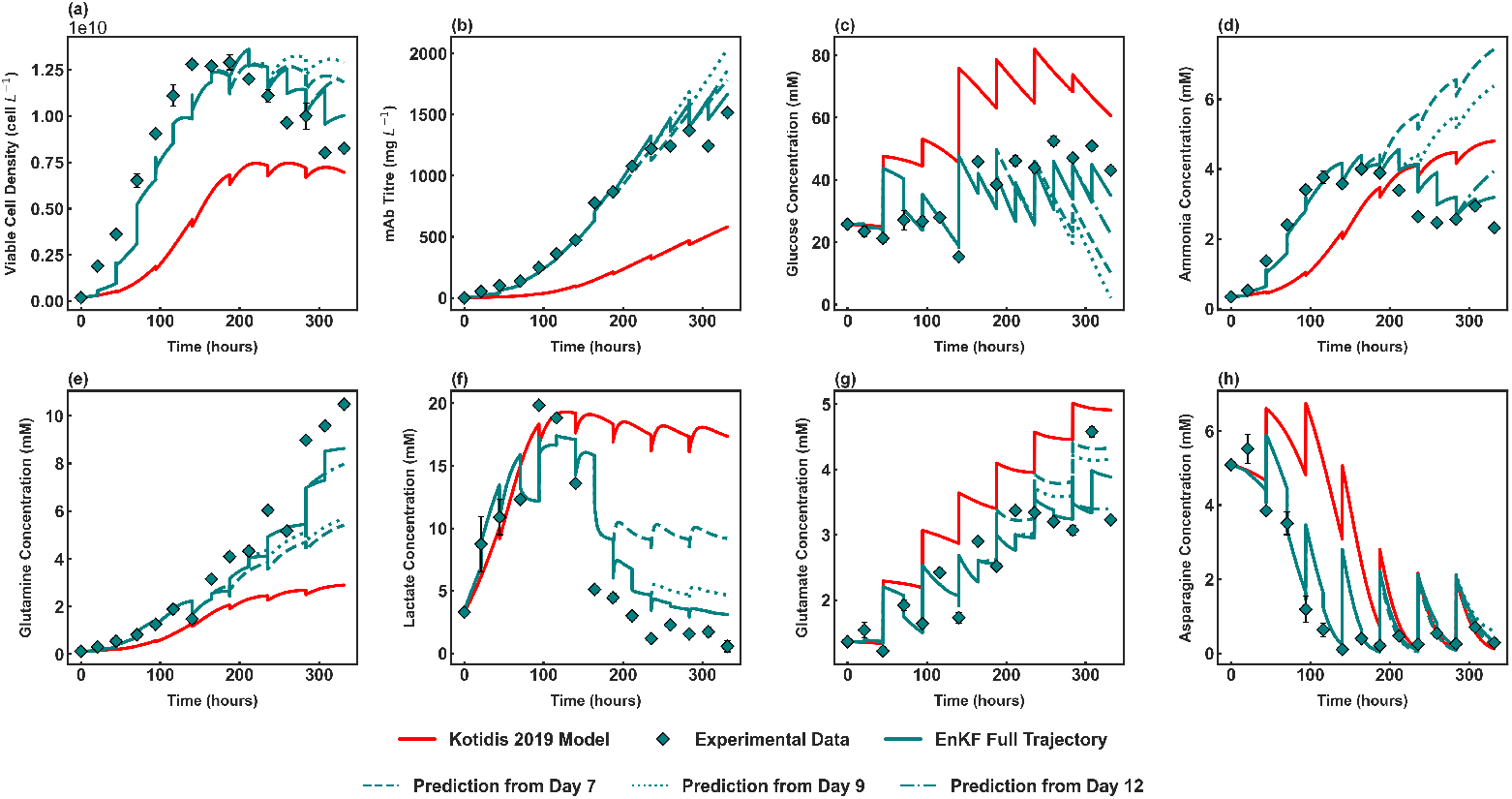
Long term EnKF predictions of Cell Line A Bioreactor at 32 °C initiated from Day 7 (dash), Day 9 (dotted), and Day 12 (dashed-dot).

**Figure S4:**
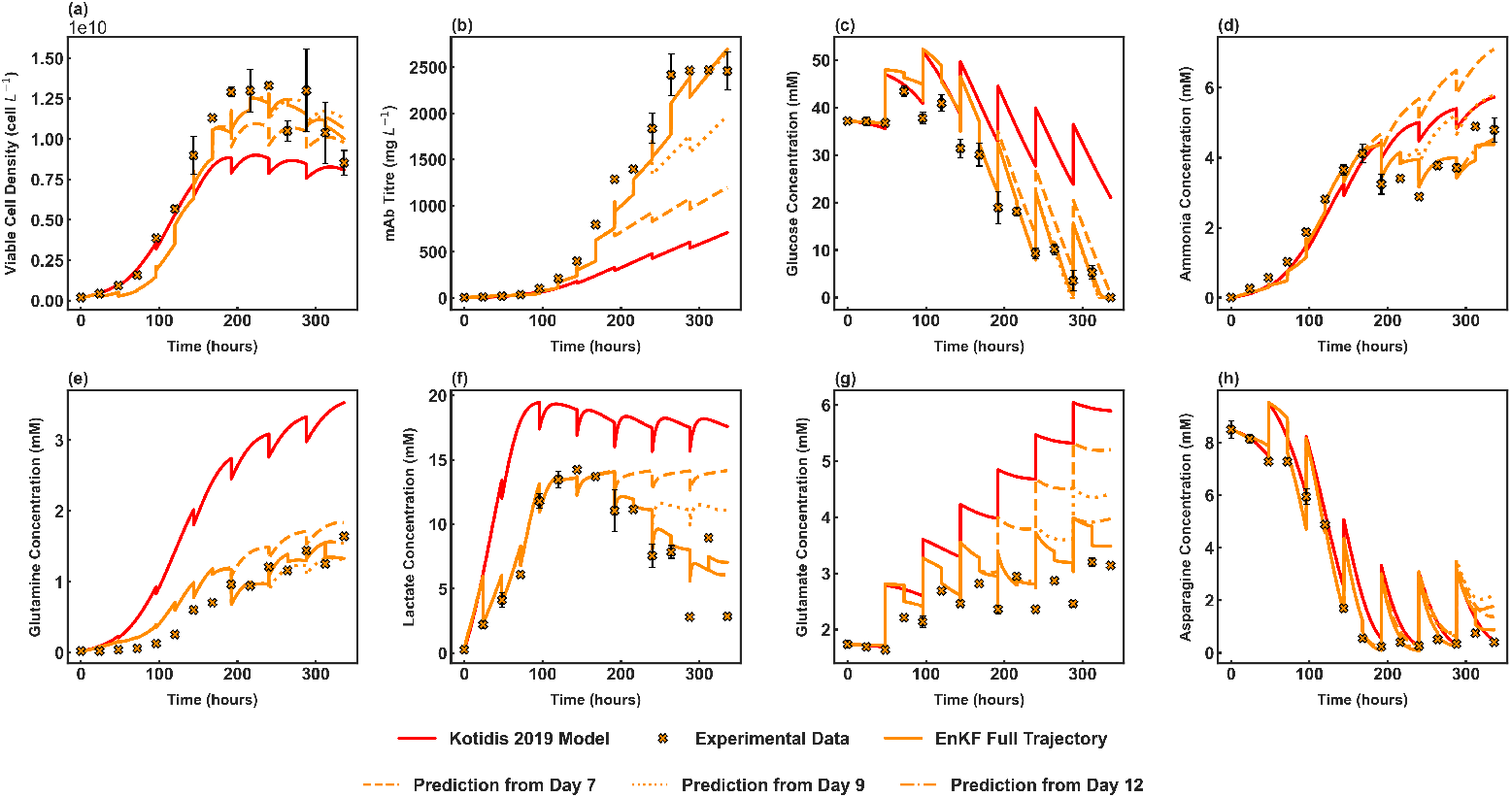
Long term EnKF predictions of Cell Line B Feed C initiated from Day 7 (dash), Day 9 (dotted), and Day 12 (dashed-dot).

**Figure S5:**
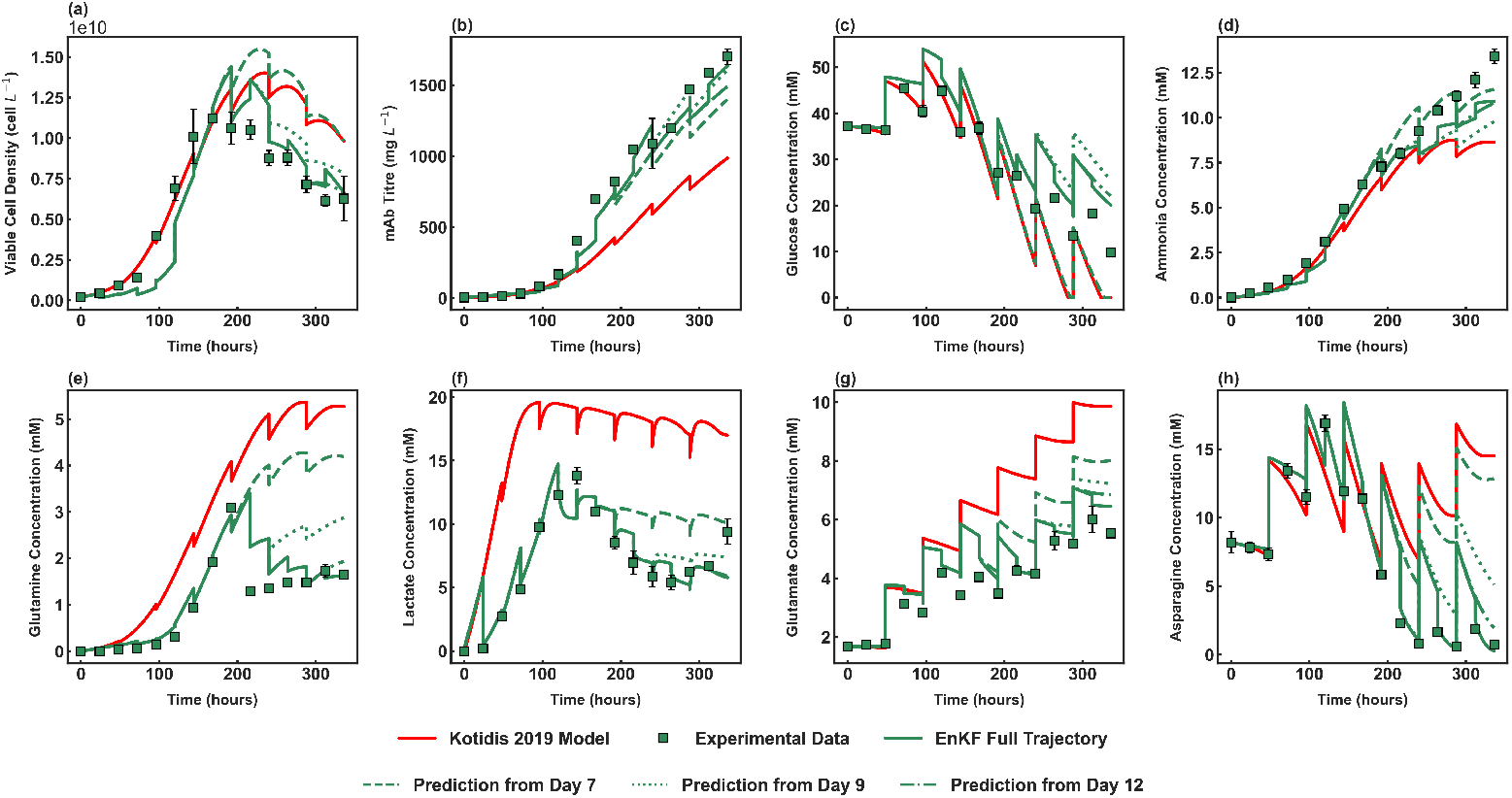
Long term EnKF predictions of Cell Line B Feed U initiated from Day 7 (dash), Day 9 (dotted), and Day 12 (dashed-dot).

